# Sex-Specific Effects of Psychedelic Drug Exposure on Central Amygdala Reactivity and Behavioral Responding

**DOI:** 10.1101/2022.04.28.489882

**Authors:** DP Effinger, SG Quadir, MC Ramage, MG Cone, MA Herman

## Abstract

Psilocybin, and its active metabolite psilocin, have been shown to elicit rapid and long-lasting symptom improvements in a variety of affective psychiatric illnesses. However, the region-specific alterations underlying these therapeutic effects remain relatively unknown. The central amygdala (CeA) is a primary output region within the extended amygdala that is dysregulated in affective psychiatric disorders. Here, we measured CeA activity using the activity marker c-Fos and CeA reactivity using fiber photometry paired with an aversive air-puff stimulus. We found that psilocin administration acutely increased CeA activity in both males and females and increased stimulus specific CeA reactivity in females, but not males. In contrast, psilocin produced time-dependent decreases in reactivity in males, but not females as early as 2-days and lasting to 28-days post administration. We also measured behavioral responses to the air-puff stimulus and found sex-dependent changes in threat responding but not exploratory behavior or general locomotion. Repeated presentations of the auditory component of the air-puff were also performed and sex-specific effects of psilocin on CeA reactivity to the auditory-alone stimulus were also observed. This study provides new evidence that a single dose of psilocin produces sex-specific, time-dependent, and enduring changes in CeA reactivity and behavioral responding to specific components of an aversive stimulus.

## INTRODUCTION

Clinical studies suggest that psilocybin, and active metabolite psilocin [1–3], produce rapid, long-lasting improvements in psychiatric disorders including depression, anxiety, and substance use disorder [4–12]. Sustained improvements in symptoms were seen as long as 4 years following administration [13]. Research on psychedelic compounds has focused primarily on human imaging studies and receptor pharmacology. Human fMRI studies have shown that psychedelics produce robust alterations in network connectivity [8, 14–19].

Receptor studies have demonstrated that these compounds promote synaptogenesis [20–22] and identified the serotonin 5-hydroxytryptamine (5-HT)_2A_ receptor to be instrumental in mediating the hallucinogenic actions of psychedelic compounds including psilocybin and lysergic acid diethylamide (LSD) [16, 23–28]. Despite the observed clinical outcomes in psychedelic studies, the underlying neural correlates of these therapeutic effects remain relatively unknown.

The central amygdala (CeA) is a primary output region within the extended amygdala that receives input regarding internal and external arousal state and coordinates appropriate behavioral responses [29, 30]. CeA dysregulation has been implicated in anxiety, depression, and PTSD [29, 31–36]. Preclinical studies demonstrate that the psychedelic compounds LSD and 2,5-dimethoxy-4-iodoamphetamine (DOI) increase Fos expression in the amygdala [37, 38] and that systemic and local DOI administration in the amygdala promotes suppression of fear responses [38]. Psilocybin produces alterations in reactivity and connectivity of the amygdala that correlate with positive therapeutic outcome [12, 15, 17, 39–41]. One clinical imaging study reported significantly decreased amygdala reactivity 1-week following psilocybin administration, with lasting changes in affect at one-month follow-up [41].

The current study utilized a preclinical approach to determine how the active compound, psilocin, alters CeA activity, reactivity, and behavioral responding to an aversive stimulus in male and female Sprague Dawley rats. Psilocin was used as it is the active metabolite of psilocybin in the central nervous system (CNS), mediating effects of psilocybin at 5-HT_2AR_ [2, 28]. We focused on the CeA as it’s the primary output region of the amygdala and as amygdala hyperreactivity is associated with many psychiatric disorders that psilocybin shows promise in treating [42–45]. We first measured expression of c-Fos, an immediate early gene that is upregulated following neuronal activity, in the CeA of male and female rats administered psilocin (2 mg/kg) or vehicle. Next, we measured CeA reactivity using an air-puff stimulus involving a burst of air directed towards the face. The air-puff stimulus has been validated as an aversive stimulus in passive avoidance tests [46] and, due to known CeA involvement in pain modulation [47, 48], it was essential to utilize a stimulus that would evoke a response without the addition of potential confounding effects of pain. Utilizing fiber photometry in conjunction with the aversive air-puff stimulus, we were able to demonstrate a robust and reliable CeA reactivity response. We then used this model to determine how administration of psilocin altered CeA reactivity during acute exposure and at more prolonged timepoints. We observed sex-specific effects of psilocin on CeA activity as measured by c-Fos expression and in-vivo reactivity and behavioral responding. Specifically, we found that psilocin elicited acute increases in reactivity in females, but not males. Additionally, we found long-term decreases in CeA reactivity in males that were not seen in females or in the vehicle control. To test whether these effects would contribute to the development of a conditioned response to the sound of the air-puff, additional assays were performed utilizing an auditory-only air-puff stimulus, wherein an air-puff was administered with the nozzle facing outside of the box to isolate the auditory component of the air-puff. In doing so, we show differential reactivity of the CeA suggesting that the observed changes were stimulus-specific and dynamic in nature. Finally, we found that decrease in reactivity in males were primarily driven by animals that exhibited darting vs, freezing behavior in response to the air puff stimulus.

## METHODS

### Animals/Stereotaxic Surgery

Adult male and female Sprague Dawley rats (200-400 g, ~7-weeks old at arrival, Envigo, Indianapolis, IN) were group-housed in a humidity- and temperature-controlled (22°C) vivarium on a 12hr light/dark cycle with *ad libitum* access to food and water. Bilateral infusions of pGP-AAV-syn-jGCaMP7f-WPRE (Addgene plasmid# 104488-AAV9) were performed at a rate of 100nl/minute into the CeA (AP:-2.0mm; ML:+/− 3.9mm; DV:-8.0mm). Directly following injection, Doric 200um NA 0.37 silica optic fiber cannulas with a 9mm tip were inserted at DV −7.7mm and secured in place using Metabond (Parkell, Brentwood, NY).

Following intracranial surgery at ~8 weeks of age, animals were single-housed in flat lid cages with waterspout access, and food bowls to avoid damage to implants.

### Drug Administration

On the day of injection, animals received a subcutaneous (s.c.) injection of either vehicle (0.9% Saline/2% glacial acetic acid) or a 2 mg/kg psilocin (Cayman Chemical, graciously provided by Dr. Bryan Roth) dissolved in 2% glacial acetic acid, as previously described [49]. Solutions were prepared fresh on the day of testing and were stored at 4°C for 30min prior to first injection. Injections were given at a 1mg/ml volume 40min prior to the beginning of fiber photometry recordings. In the c-Fos cohort, injections were performed 2hr prior to perfusion. For this study, a dose of 2 mg/kg was chosen as previous work has shown that administration of psilocin at 2 mg/kg in rats produced increases in BOLD signaling in the amygdala, an effect not seen when utilizing a lower dose [50].

### Fiber Photometry Recording

All animals were ~11 weeks of age at the beginning of testing. Animals were habituated to testing room for 1 hour prior to recording sessions. Rats were individually placed in a clear plexiglass chamber [50cm (l)×50cm(w)×38cm(h)] in a red-light sound-attenuated behavioral cabinet and allowed to acclimate for 10min. Fiber placement and GCaMP injection were performed bilaterally, however, recordings were collected unilaterally. Baseline recordings were taken from both hemispheres and analyzed to determine which hemisphere to use for the remainder of the experiment. Ca^2+^ signals from subjects, with 200μm optic fibers placed .3mm above the injection site in the CeA, were recorded using a TDT RZ5 real time processor equipped with the TDT Synapse program (Tucker-Davis Technologies, Alachua, FL). For each recording, jGCaMP7f was excited using a 465nm calcium-dependent signal and a 405nm signal was used as an isosbestic control. MATLAB script [51] was then used to analyze raw signal. Changes in fluorescence as a function of baseline (ΔF/F) showing fluctuations in fluorescence compared to the overall baseline fluorescence throughout the session were calculated through smoothing isosbestic control channel signal, scaling the isosbestic signal by regressing it onto smoothed GCaMP signal, and then generating a predicted 405nm from the regression to remove calcium-independent signaling from the raw GCaMP signal and control for artifact from movement, photo-bleaching, and fiber bending artifacts. Baseline fluorescence was calculated through a least-squares linear fitting of the 405nm isosbestic control to the 465nm GCaMP signal to create a resting GCaMP signal over the entire session not containing event-dependent fluctuations in GCaMP signal [52]. Peri-event plots were then generated by averaging changes in fluorescence (ΔF/F) across two different air-puff trials to generate a mean change in reactivity [51]. For all photometry experiments, the same male experimenter handled animals and recording. Additionally, female animals were not checked for estrus cycle to avoid potential confounding effects of an added stressor. All subjects were video recorded during stimulus presentations and behavior was scored using the Behavioral Observation Research Interactive Software (BORIS) program [53]. Behavioral analysis was performed by an experimenter blind to the experimental status of each animal.

### Air-Puff Stimulus

An aversive air-puff stimulus was designed using house air supply (85psi) controlled by a Parker solenoid (Part #003-0868-900), powered by a custom made 12V electrical circuit box connected to the TDT RZ5 system to trigger 500ms openings of the solenoid and simultaneous timestamps through the Synapse program. Each session consisted of a 10min habituation period before recording sessions that consisted of two air-puff administrations with a 5min inter-stimulus interval (ISI) between them. Air-puffs were directed at the face and within close proximity of the animal to maintain similar conditions and level of aversion to the stimulus between subjects.

### Perfusion/Tissue Collection

Rats were transcardially perfused with phosphate buffered saline (PBS) followed by 4% paraformaldehyde (w*/v*). Brains were extracted, stored in 4% paraformaldehyde for 24h, then transferred to 30% sucrose (*w/v*) and stored at 4°C until sectioning. Using a freezing microtome, tissue was sliced into 40μm thick sections and stored in cryoprotectant (30%*v/v* ethylene glycol + 30% *w/v* sucrose in phosphate buffered saline) at 4°C until use.

### Immunofluorescence Staining

For site verification on the fiber photometry cohorts, 6 sections/subject (anterior-posterior (AP) coordinates −1.75 to −2.75) from bregma were washed in phosphate buffered saline (PBS) prior to 1hr blocking in 5% normal donkey serum + 0.3% Triton X in PBS. Sections were then incubated in chicken anti-GFP (Abcam 13970; 1:1000) in 0.5% normal donkey serum/0.3% Triton X/PBS overnight at 4°C. Next, samples were triple-washed and incubated in blocking solution (1hr) followed by goat anti-chicken 488 (Jackson 703-545-155; 1:700) in 0.5% normal donkey serum/0.3% Triton X/PBS for 2hr at room temperature.

For the c-Fos cohort, sections (AP coordinates −1.92mm to −2.16mm) were washed in PBS, incubated in 50% methanol/PBS solution for 30min, washed in 3% hydrogen peroxide/PBS for 5min, placed in blocking solution (0.3%Triton X-100; Thermo Fisher), and then 1% bovine serum albumin (BSA; Sigma) for 1hr, at room temperature (RT). Slices were then incubated with rabbit anti-*cFos* (1:3000, Millipore Sigma; ABE457) for 24hr at 4°C. Next, slices were washed with 0.1% Tween-20 in tris-buffered saline (TNT) before being transferred into TNB blocking buffer (Perkin-Elmer FP1012) for 30min. After blocking, slices were incubated in goat anti-rabbit horseradish peroxidase (HRP; 1:200, Abcam ab6721) for 2hr followed by another round of TNT washes. Finally, slices were incubated in tyramide conjugated fluorescein (1:50) in TSA amplification diluent (Akoya Biosciences, NEL741001KT) for 10min at RT. Slices were washed with TNT buffer, mounted, cover slipped with Vectashield ^®^ HardSet™ Antifade Mounting Medium with DAPI (H1500, Vector Laboratories, Burlingame, CA) and stored at 4°C before being imaged with the Keyence BZ-X800 fluorescence microscope. Images were then analyzed by an experimenter blinded to the animal’s condition. Images were taken at 20X and the BZ-X800 Analyzer program was used to stitch images together. Randomly selected hemispheres were used, and the area of interest was outlined and manually counted using the ImageJ [54] multi-point counter tool. Cell counts were averaged per animal.

### Statistical Analysis

Raw data from the TDT RZ5 machine were imported and processed for artifact removal, down sampling, and detrending using MATLAB code [51]. Changes in fluorescence (ΔF/F) were then calculated and collected 5s before and 10s following air-puff administration. The mean of two traces were averaged together for each subject and combined using custom MATLAB script (https://doi.org/10.5061/dryad.pnvx0k6q9) to sort individual data into groups. Group 465nm signal data were then converted into 500ms bins for analysis to quantify differences between experimental conditions. All binned trace plots were normalized by subtracting the mean of the signal pre-air-puff from each time point in order to correct for differences in basal activity prior to air puff administration. All comparisons were made using time bins post air puff, corresponding to 0-20 on the x-axis of binned analysis plots. All statistical analyses were conducted using GraphPad Prism 7. Five subjects were removed from the study due to lack of signal/cannula removal. Repeated-measures one-way analysis of variance (ANOVA) with Dunnett’s post hoc test was used to determine differences between baseline, injection day, and follow-up time points within each group in the air-puff condition. Two-way ANOVAs with Šidák’s post hoc tests were used to compare time bins between groups and time points. Paired t-tests were used to determine differences within groups between the peak point between the intial and final auditory only recording days. Fisher’s exact tests were used to compare percentages in darting behavior. Mixed-effects two-way ANOVA were used to compare time spent walking and rearing. For all binned trace data plots, a bootstrapping confidence interval (CI) procedure (99% CI, 1000 iterations) [55] was conducted to calculate a new ‘mean”ΔF/F by randomly resampling rom subject mean ΔF/F with replacement corresponding to the number of samples. The *n* for all analyses corresponded to the number of subjects in each group. This process was repeated 1000 times to create a nonparametric distribution of population mean simulations for each time point within the window. A statistically significant increase (>0%) in Ca^2+^ transients following the air puff administration was defined as those data points wherein the lower bound of the 99% CI was >0. [55–57] This was signified on graphs by color matched lines above the trace bin plots. All data including Matlab structures and scripts can be found at https://doi.org/10.5061/dryad.pnvx0k6q9.

## RESULTS

### Acute Changes in CeA Reactivity in Response to an Aversive Air-Puff Stimulus

At ~8 weeks, male and female Sprague Dawley rats received bilateral injection of pGP-AAV-syn-jGCaMP7f-WPRE into the CeA immediately followed by installation of fiber optic cannulae above the injection site (**Fig. 1A-B**). Following 3-weeks for virus transduction, animals underwent a series of fiber photometry experiments testing the effects of psilocin on fluctuations in calcium dynamics within the CeA time-locked to an aversive air-puff stimulus (**Fig. 1A**). For both baseline and injection recording sessions, each animal received two air-puffs with an ISI of 5min and trace plots were generated showing change in fluorescence as a function of baseline (ΔF/F). Traces were analyzed 5s before and 10s following administration of air-puff (**Fig. 1C**). Following injection of the drug, animals were not assessed for any behavioral indices of hallucination (i.e. head twitch, wetback shake).

**Figure 1.**
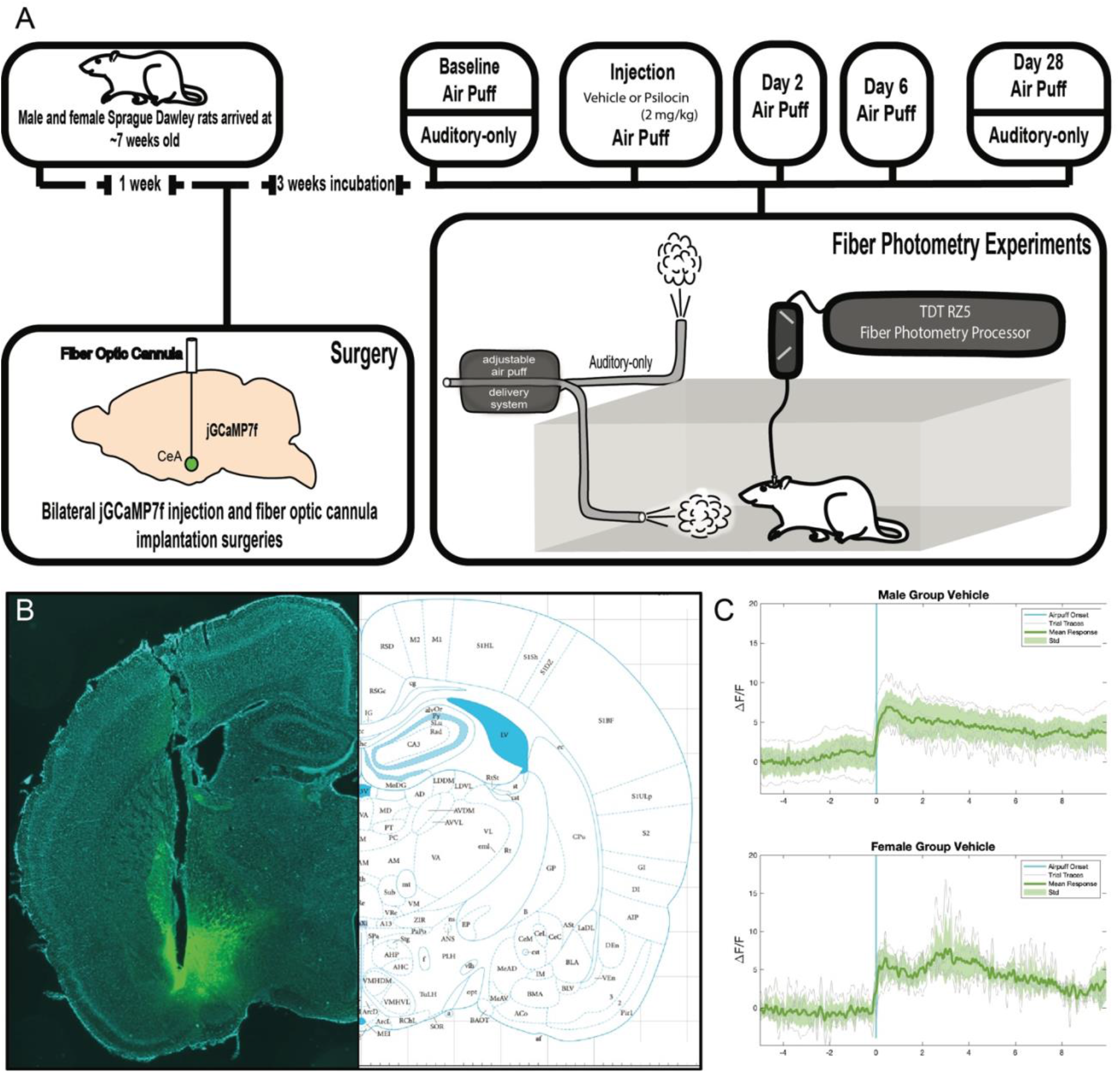
(**A**) Experimental timeline. Animals arrived at 6-7 weeks old and were allowed to habituate for 1-week prior to surgeries. Bilateral injection of calcium sensor (jGCaMP7f) and implantation of fiber optic cannulas were conducted. After 3 weeks for viral transduction, animals underwent a series of fiber photometry experiments. (**B**) Representative coronal section showing fiber placement and GCaMP expression. (**C**) Mean air-puff traces showing consistency of CeA signal in response to the air-puff stimulus.

### Sub-Region-Specific Differences in CeA Basal Activity Following Psilocin Administration

To assess the effects of psilocin on basal activity in the CeA, male and female Sprague Dawley rats were injected with psilocin (2 mg/kg, s.c.) or vehicle (1 ml/kg, s.c.) 2hr prior to transcardial perfusion.

Immunohistochemistry (IHC) assays were performed using CeA tissue collected within the anterior-posterior bregma range of −1.92mm to −2.16mm. In females, there was a significantly increased c-Fos expression in the psilocin group as compared to vehicle across regions (2-way ANOVA: F_Interaction_(2,24)=1.597, *p*=0.22; F_Region_(1,24)=30.84, p<0.0001; F_injection_(1,24)=9.149 *p*=0.006; **Fig.2A-B**) with greater increases in the capsular division of the central amygdala (CeC) (Šidák’s: *p*=0.02). In males, there was also significantly increased c-Fos expression in the psilocin group across regions (2-way ANOVA: F_Interaction_(2,27)=1.241, *p*=0.30; F_Region_(2,27)=12.02, p=0.0002; F_injection_(1,27)=5.543, *p*=0.02; **Fig.2C-D**).

**Figure 2.**
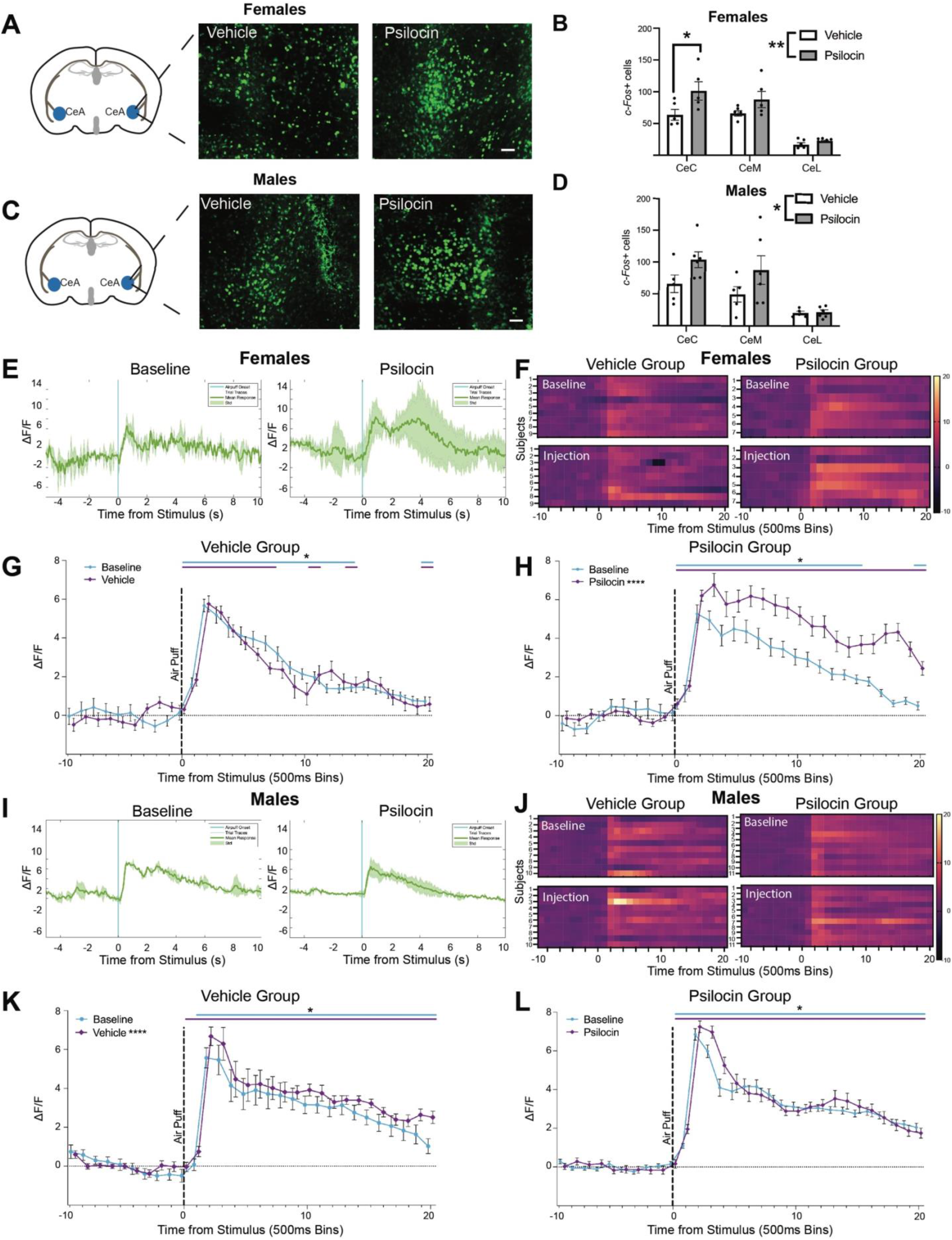
Acute Changes in CeA Reactivity in Response to an Aversive Air-Puff Stimulus Following Psilocin Administration. **(A)** Females: representative images showing c-Fos+ cells tagged with a green fluorescence protein (GFP) in the CeA. Scale bar = 100 **μm. (B)** Female vehicle vs. psilocin: histogram showing the number of c-Fos+ cells in each subregion of the CeA. CeC = capsular CeA, CeM =. Medial CeA, CeL = lateral CeA. Data points correspond to the mean of 2-4 hemispheres for each individual subject. **(C)** Males: representative images showing c-Fos+ cells tagged with a green fluorescence protein (GFP) in the CeA. Scale bar = 100 **μm. (D)** Male vehicle vs. psilocin: histogram showing the number of c-Fos+ cells in each subregion of the CeA. CeC = capsular CeA, CeM =. Medial CeA, CeL = lateral CeA. Data points correspond to the mean of 2-4 hemispheres for each individual subject. **(E)** Female psilocin group: representative raw ΔF/F traces showing an individual at baseline and while on psilocin. Blue line = air-puff onset, Grey lines = individual traces, Green line = mean trace, Std= standard deviation **(F)** Females: Heatmaps comparing baseline to injection. Each row represents an individual subjects mean trace. (**G)** Female vehicle group: air-puff trace plots of changes in CeA fluorescence following exposure to a 500ms air-puff at 85 psi. Data points represent group averages within 500ms binned window +/− S.E.M. **(H)** Female psilocin group: air-puff trace plots of changes in CeA fluorescence following exposure to a 500ms air-puff at 85 psi. Data points represent group averages within 500ms binned window +/− S.E.M. **(I)** Male psilocin group: representative raw ΔF/F traces showing an individual at baseline and while on psilocin. Blue line = air-puff onset, Grey lines = individual traces, Green line = mean trace, Std= standard deviation **(J)** Males: Heatmaps comparing baseline to injection. Each row represents an individual subjects mean trace. **(K)** Male vehicle group: air-puff trace plots of changes in CeA fluorescence following exposure to a 500ms air-puff at 85 psi. Data points represent group averages within 500ms binned window +/− S.E.M. **(L)** Male psilocin group: air-puff trace plots of changes in CeA fluorescence following exposure to a 500ms air-puff at 85 psi. Data points represent group averages within 500ms binned window +/− *Prolonged Changes in CeA Reactivity in Response to an Aversive Air-Puff Stimulus Following Psilocin Administration*

### Acute Changes in CeA Reactivity in Response to an Aversive Air-Puff Stimulus Following Psilocin Administration

To test the acute effects of psilocin on CeA reactivity, we measured changes in CeA reactivity during time-locked stimuli presentation at baseline and 40min post administration of vehicle or psilocin (2 mg/kg, s.c.). Following a 10min habituation period, air-puffs were delivered as described above. Mean traces were taken from two separate air-puff responses within a session (**Fig. 2E, 2I**). These traces were then collapsed into 500ms bins for statistical analysis (**Fig. 2G, 2H, 2K, 2L**). A repeated measures one-way analysis of variance (ANOVA) was used to determine an effect of all the different timepoints (baseline, injection, 2-day, 6-day, and 28-day) within each group. We utilized a bootstrapping 99% confidence interval (CI) procedure to estimate population mean ΔF/F during the 10 time bins (5s) prior and 20 time bins (10s) following air puff administration. A significant increase in ΔF/F from 0 to 20 time bins (0-10s) following air puff was determined whenever the lower bound of the 99% CI was >0. These points of statistical significance are shown above binned trace ΔF/F plots and are colored coded to match the corresponding time bin trace. In the female vehicle control group (n=9), there was a main effect of time points (F_(1.558,29.61)_=29.32, p<0.0001). However, there were no changes between baseline and injection (*p*=0.36; **Fig.2F-G**). In the female psilocin group (n=7), there was a significant effect of time points (F_(3.178,60.38)_=50.25, p<0.0001) with a significant increase between baseline and injection (*p*<0.0001; **Fig.2F, 2H**). In the male vehicle control group (n=10), there was main effect of time points (F_(1.797,34.14)_=14.48) with an increase in reactivity between baseline and injection (*p*<0.0001; **Fig.2J-K**), though this effect appears to be primarily driven by a single subject as opposed to uniform increases across all subjects (**Fig.2J**). In the male psilocin group (n=11), there was a main effect of time points (F_(2.163,41.10)_=63.78) with no significant difference between baseline and injection (*p*=0.47; **Fig.2J, 2L**).

To assess long-term alterations in CeA reactivity following a single dose of psilocin, fiber photometry recordings were performed on days 2, 6, and 28 post-administration of vehicle or psilocin. On each follow-up recording day, animals received two air-puffs with a 5-minute ISI. In the vehicle control females, there was an effect of time points (F_(1.558,29.61)_=29.32, *p*<0.0001; **Fig.3B,3C**), with significant increases in reactivity seen at the 2-day (*p*<0.0001) and 28-day follow-up (*p*=0.0005) compared to baseline. In the female psilocin group (**Fig. 3A, 3D**), there was an effect of time points (F_(3.178,60.38)_=50.25, *p*<0.0001; **Fig.3B,3D**), with significant increases in reactivity at the 2-day (*p*=0.02), 6-day (*p*<0.0001), and 28-day follow-up (*p*=0.04).

**Figure 3.**
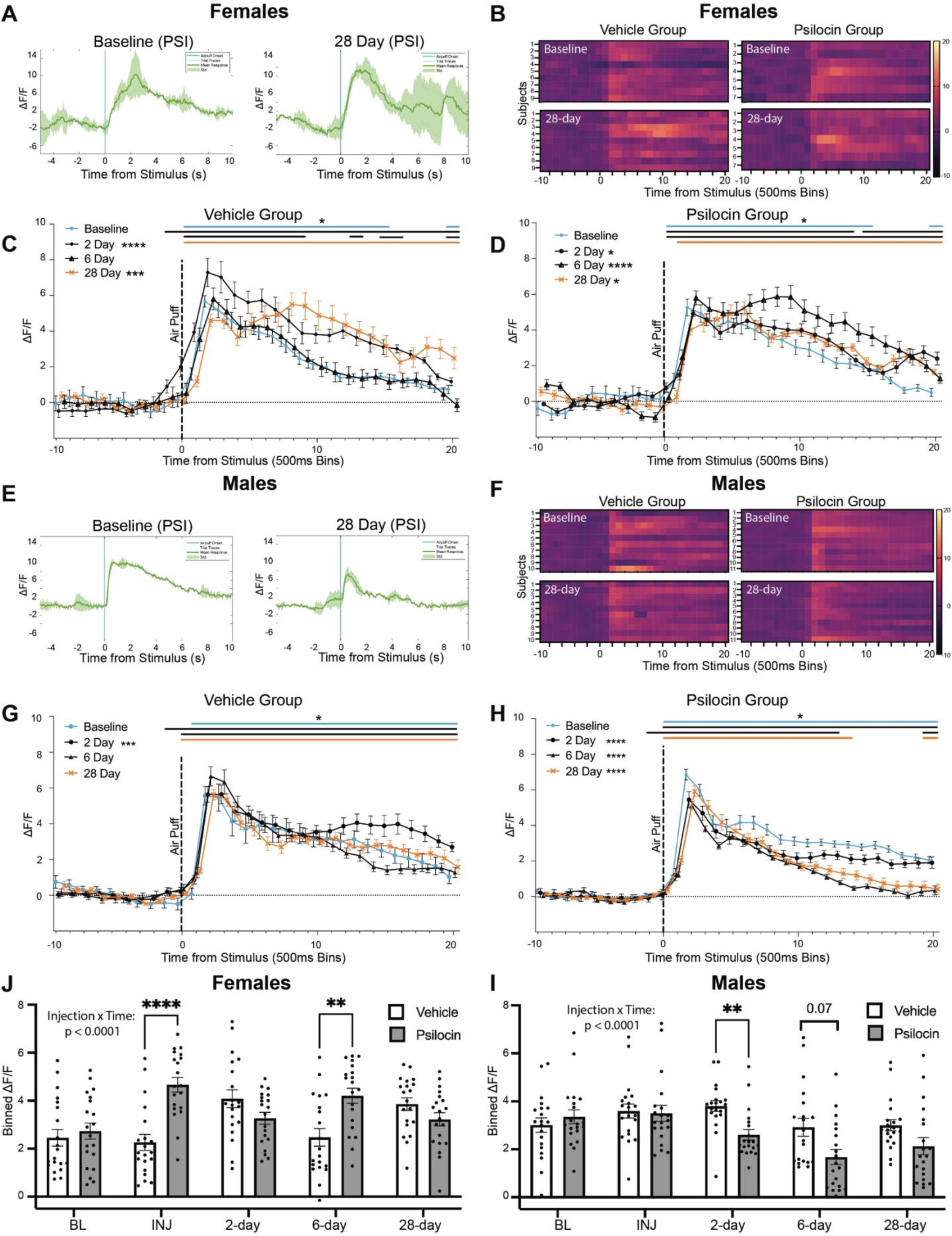
Prolonged Changes in CeA Reactivity in Response to an Aversive Air-Puff Stimulus Following Psilocin Administration. **(A)** Female psilocin group: representative raw ΔF/F traces showing an individual at baseline and at the 28-day follow-up. Blue line = air-puff onset, Grey lines = individual traces, Green line = mean trace, Std= standard deviation **(B)** Females: Heatmaps comparing baseline to the 28-day follow up. Each row represents an individual subjects mean trace. **(C)** Female vehicle group: air-puff trace plots of changes in CeA fluorescence following exposure to a 500ms air-puff at 85 psi. Data points represent group averages within 500ms binned window +/− S.E.M. **(D)** Female psilocin group: air-puff trace plots of changes in CeA fluorescence following exposure to a 500ms air-puff at 85 psi. Data points represent group averages within 500ms binned window +/− S.E.M. **(E)** Male psilocin group: representative raw ΔF/F traces showing an individual at baseline and while on psilocin. Blue line = air-puff onset, Grey lines = individual traces, Green line = mean trace, Std= standard deviation **(F)** Males: Heatmaps comparing baseline to the 28-day follow up. Each row represents an individual subjects mean trace. **(G)** Male vehicle group: air-puff trace plots of changes in CeA fluorescence following exposure to a 500ms air-puff at 85 psi. Data points represent group averages within 500ms binned window +/− S.E.M. **(H)** Male psilocin group: air-puff trace plots of changes in CeA fluorescence following exposure to a 500ms air-puff at 85 psi. Data points represent group averages within 500ms binned window. **(I)** Females: Summary histogram comparing the mean of all 20 time bins following air-puff administration. Each data point represents an individual time bin within that condition collapsed across all subjects. +/− S.E.M. **(J)** Males: Summary histogram comparing the mean of all 20 time bins following air-puff administration +/− S.E.M. Each data point represents an individual time bin within that condition collapsed across all subjects. +/− S.E.M. In each trace bin plot panel, a significant increase in ΔF/F was determined whenever the lower bound of the 99% CI was >0. These points of statistical significance are shown as colored lines above each ΔF/F curve with colors corresponding to the respective binned traces with a * above the lines. \**p*<0.05, \*\**p*<0.01, \*\*\**p*<0.001, \*\*\*\**p*<0.0001.

In the male vehicle group, there was an effect of time point (F_(1.797,34.14)_=14.48, *p*<0.0001; **Fig.3F,3G**), with a significant increase in reactivity at the 2-day follow-up (*p*<0.0001). In the male psilocin group, there was a significant effect of time point (F_(2.163,41.10)_=63.78, *p*<0.0001; **Fig.3F,3H**), with significant decreases in reactivity seen at the 2-day, 6-day, and 28-day follow-up (*p*<0.0001).

To compare changes in reactivity between vehicle and psilocin groups, 2-way ANOVA revealed differential effects of injection across time points between groups in females (F_Interaction_(4,152)=53.46, *p*<0.0001; F_Injection_(1,38)=2.015, *p*=0.16; F_Time_(2.343,89.04)=18.82, *p*<0.0001; **Fig.3J**) and males (F_Interaction_(4,152)=23.96, *p*<0.0001; F_Injection_(1,38)=2.310, *p*=0.13, F_Time_(2.603,98.90)=50.45, *p*<0.0001; **Fig.3I**). Šidák post hoc analysis revealed significant increases in reactivity during injection (*p*<0.0001) and at the 2-day follow-up (*p*=0.005) in the female psilocin group compared to vehicle control (**Fig.3J**). In males, there was a significant decrease in reactivity at the 2-day follow-up in the psilocin group compared to vehicle control (*p*=0.003; **Fig.3J**).

### Acute and Prolonged Behavioral Effects of Psilocin

To determine if psilocin elicited changes in locomotion and exploratory behavior, video recordings were scored for total time spent walking and time spent rearing, respectively. All behavior was scored during the 5min ITI between the first and second air-puff. Rearing was defined as standing on the two hindlegs, regardless of any balancing on the walls of the box. In females, there were no differences in overall locomotion between vehicle and psilocin groups (F_Interaction_(3,42)=0.2118, *p*=0.89; F_Injection_(1,14)=2.238, *p*=0.16; F_Time_(2.497,34.95)=0.6065, *p*=0.59; **Fig.4A**). There were also no differences in rearing between female vehicle and psilocin groups (F_Interaction_(4,56)=1.345, *p*=0.26; F_Injection_(1,14)=4.433, *p*=0.05; F_Time_(4,56)=2.486, *p*=0.05; **Fig. 4B**). In the males, there were no differences in locomotion (F_Interaction_(3,57)=0.1054, *p*=0.96; F_Injection_(1,19)=2.563, *p*=0.12; F_Time_(3,57)=1.255, *p*=0.29; **Fig.4C**) or rearing (F_Interaction_(4,76)=0.5546, *p*=0.69; F_Injection_(1, 19)=0.7576, *p*=0.39; F_Time_(4,76)=1.190, *p*=0.32; **Fig.4D**) between vehicle and psilocin groups. Collectively, no significant alterations in locomotion were seen resulting from psilocin administration (**Fig.4A, 4C**), although females appeared to display increased locomotion overall as compared to males, regardless of injection type.

**Figure 4.**
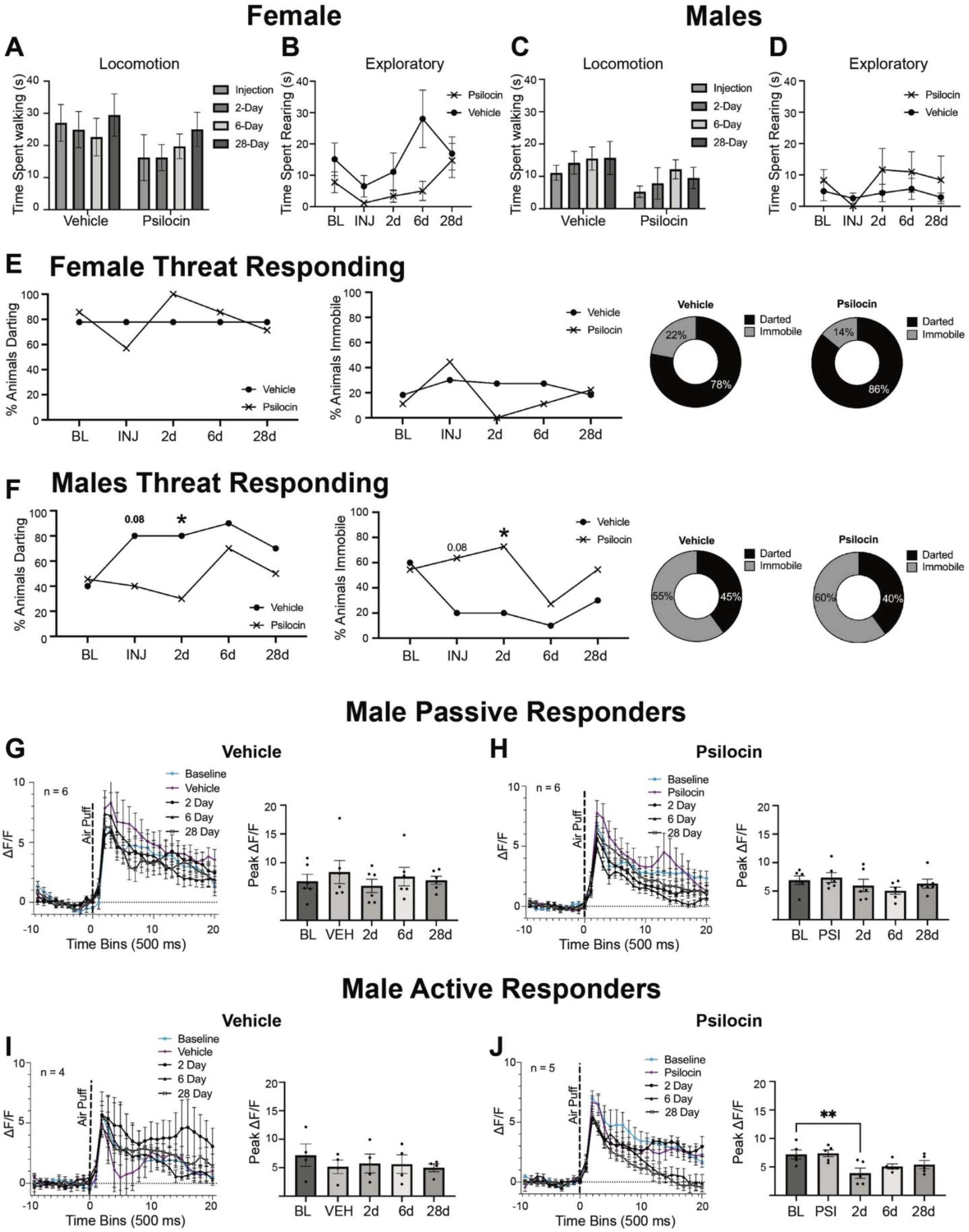
Acute and Prolonged Behavioral Effects of Psilocin. **(A)** Females: locomotion plots comparing vehicle and psilocin groups at each time point. Data points are mean time in seconds +/− SEM. **(B)** Females: time spent rearing comparing vehicle and psilocin groups at each time point. Data points are mean time in seconds +/− SEM. **(C)** Males: locomotion plots comparing vehicle and psilocin groups at each time point. Data points are mean time in seconds +/− SEM. **(D)** Males: time spent rearing comparing vehicle and psilocin groups at each time point. Data points are mean time in seconds +/− SEM. **(E)** Females threat responding behavior. First figure is showing percentage of subjects that darted following air-puff administration. Second figure is showing the percentage of animals that remained immobile following air-puff. Third and fourth figure are pie charts showing the percentage of animals that darted vs. remained immobile **(F)** Males threat responding behavior: First figure is showing percentage of subjects that darted following air-puff administration. Second figure is showing the percentage of subject that remained immobile following air-puff. Third and fourth figure are pie charts showing the percentage of animals that darted vs. remained immobile **(G)** Male vehicle passive responders (remained immobile) group: First figure shows air-puff trace plots of changes in CeA fluorescence following exposure to a 500ms air-puff at 85 psi. Data points represent group averages within 500ms binned window +/− S.E.M.; Second figure is a histogram showing differences in the mean peak ΔF/F value for each group +/− S.E.M following air-puff administration. Data points reflect each individual subject within the corresponding subgroup. **(H)** Male Psilocin passive responders (remained immobile) group: First figure shows air-puff trace plots of changes in CeA fluorescence following exposure to a 500ms air-puff at 85 psi. Data points represent group averages within 500ms binned window +/− S.E.M. Second figure is a histogram showing differences in the mean peak ΔF/F value for each group +/− S.E.M following air-puff administration. Data points reflect each individual subject within the corresponding subgroup. **(I)** Male vehicle active responders (darted) group: First figure shows air-puff trace plots of changes in CeA fluorescence following exposure to a 500ms air-puff at 85 psi. Data points represent group averages within 500ms binned window +/− S.E.M. Second figure is a histogram showing differences in the mean peak ΔF/F value for each group +/− S.E.M following air-puff administration. Data points reflect each individual subject within the corresponding subgroup. **(J)** Male Psilocin active responders (darted) group: First figure shows air-puff trace plots of changes in CeA fluorescence following exposure to a 500ms air-puff at 85 psi. Data points represent group averages within 500ms binned window +/− S.E.M. Second figure is a histogram showing differences in the mean peak ΔF/F value for each group +/− S.E.M following air-puff administration. Data points reflect each individual subject within the corresponding subgroup.

To probe potential differences in threat responding to the air-puff stimulus, we separated groups based on active or passive coping strategy. Subgroups were created containing animals that remained immobile (passive) following air-puff administration versus animals that exhibited darting behavior (active) following air-puff administration. Consistent with previous literature [58], females primarily employed an active darting response (**Fig.4E**), while active vs. passive coping behavior was more evenly split in males (**Fig.4F**). In females, there were no differences in the percentage of animals that darted or that stayed immobile between groups at any of the time points (**Fig. 4E**). In contrast, the male psilocin group displayed nonsignificant increases in darting and reductions in immobility in the vehicle control group that were not seen in the psilocin group under injection conditions (Fisher’s exact: *p*=0.08; **Fig. 4F**). At the 2-day follow-up, there was a significantly higher percentage of darting in the vehicle control group compared to the psilocin group (*p*=0.03; **Fig.4F, left**) and a significantly higher percentage immobile in the psilocin group (Fisher’s exact: *p*=0.03; **Fig.4F, right**). Though differences were not significant at every time point, psilocin seemed to prevent a trending increase in darting behavior seen in the male vehicle control group and increase in overall immobility in male but not female rats (**Fig.4F**).

To explore the CeA reactivity correlates to the observed behavioral differences in threat responding, fiber photometry traces were split into the corresponding active vs. passive responder groups. Since females primarily employed active coping strategies that were not affected by psilocin, we only examined CeA reactivity between the male active and passive responders. To assess changes in the amplitude of CeA response to the stimulus, comparisons looking at the peak point, or highest ΔF/F value following air-puff, were made. In the male passive responder, there were no significant differences in peak point in the vehicle (F(4,20)=1.019, *p*=0.42; **Fig.4G**) or psilocin group (F(4,20)=1.278, *p*=0.31; **Fig.4H**). In the active responder vehicle males, no differences were seen in peak point compared to baseline (F(4, 12)=1.959, *p*=0.16; **Fig.4I**). However, in the active responder psilocin males, there was a main effect of injection (F(4,16)=6.398, *p*=0.002; **Fig.4J**) with a significant decrease in peak point at day 2 compared to baseline (*p*=0.003), suggesting that psilocin-induced decreases in the amplitude of CeA reactivity were more pronounced in males employing an active vs. passive threat response at baseline.

### Changes in CeA Reactivity in Response to an Auditory Stimulus Following Psilocin Administration

To test whether the animals developed a conditioned response to the tone associated with the air-puff stimulus, and whether psilocin treatment altered the response to the auditory stimulus alone, two separate auditory-only sessions were conducted immediately following baseline recordings and again at the 28-day recording session. These auditory only sessions followed the same parameters as the air-puff with the exceptions being that the ISI was reduced to 3min and the air-puff apparatus was facing outside of the box to maintain the same auditory tone without the physical sensation of the air-puff.

In vehicle control females (**Fig.5A**), there was a significant decrease in peak point reactivity from the initial to the final auditory condition (t(8)=2.806, *p*=0.02; **Fig.5B-D**). In psilocin females (**Fig.5E**), there were no significant differences between the initial and final auditory session in peak point (t(6)=0.5642, *p*=0.59; **Fig.5F-H)**. In the vehicle males (**Fig.6A**), there weren’t any differences in peak point (t(9)=0.8811, *p*=0.40; **Fig. 6B-D**) between the initial and final auditory stimulus. Similarly, in the psilocin-treated males (**Fig.6E**), there was no difference in peak point from the initial to the final auditory session (t(10)=0.5758, *p*=0.57; **Fig.6F-H**).

**Figure 5.**
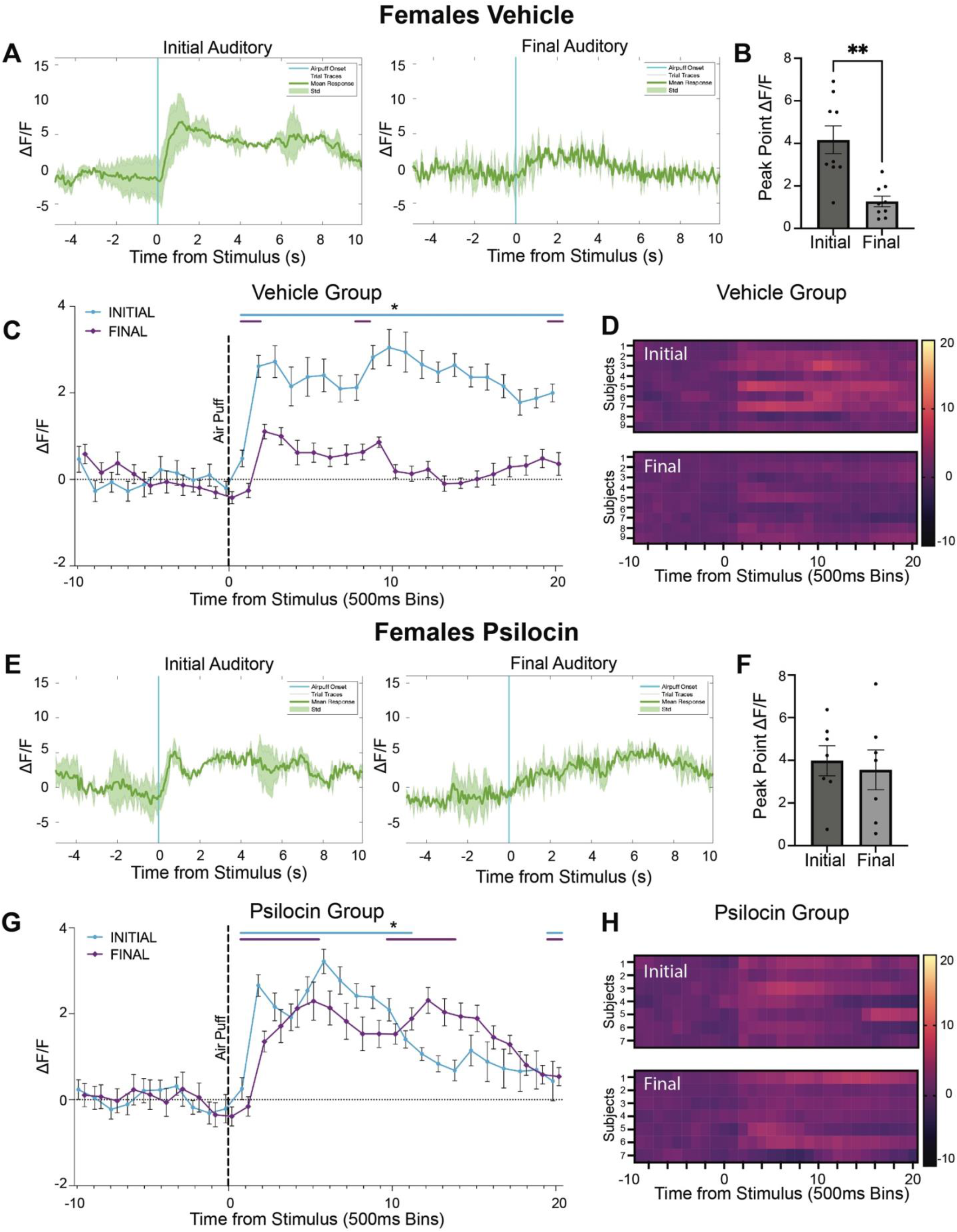
Changes in CeA Reactivity in Response to an Auditory Stimulus Following Psilocin Administration in Female Subjects. **(A)** Female vehicle group: representative mean ΔF/F traces showing an individual at initial auditory compared to final auditory. Blue line = air-puff onset, Grey lines = individual traces, Green line = mean trace, Std= standard deviation **(B)** Female vehicle group: peak point of binned mean ΔF/F traces compared between initial and final auditory recordings. Data points represent each individual subject’s peak ΔF/F value +/− SEM. **(C)** Female vehicle group: air-puff trace plots of changes in CeA fluorescence following exposure to a 500ms air-puff at 85 psi. Data points represent group averages within 500ms binned window +/− S.E.M **(D)** Female vehicle group: Heatmaps comparing initial to final auditory. Each row represents an individual subjects mean trace. **(E)** Female psilocin group: representative mean ΔF/F traces showing an individual at initial auditory compared to final auditory. Blue line = air-puff onset, Grey lines = individual traces, Green line = mean trace, Std= standard deviation **(F)** Female psilocin group: peak point of binned mean ΔF/F traces compared between initial and final auditory recordings. Data points represent individual subject’s peak ΔF/F value +/− SEM. **(G)** Female psilocin group: air-puff trace plots of changes in CeA fluorescence following exposure to a 500ms air-puff at 85 psi. Data points represent group averages within 500ms binned window +/− S.E.M. **(H)** Female psilocin group: Heatmaps comparing initial to final auditory. Each row represents an individual subjects mean trace. In each trace bin plot panel, a significant increase in ΔF/F was determined whenever the lower bound of the 99% CI was >0. These points of statistical significance are shown as colored lines above each ΔF/F curve with colors corresponding to the respective binned traces with a * above the lines. \**p*<0.05, \*\**p*<0.01, \*\*\**p*<0.001, \*\*\*\**p*<0.0001.

**Figure 6.**
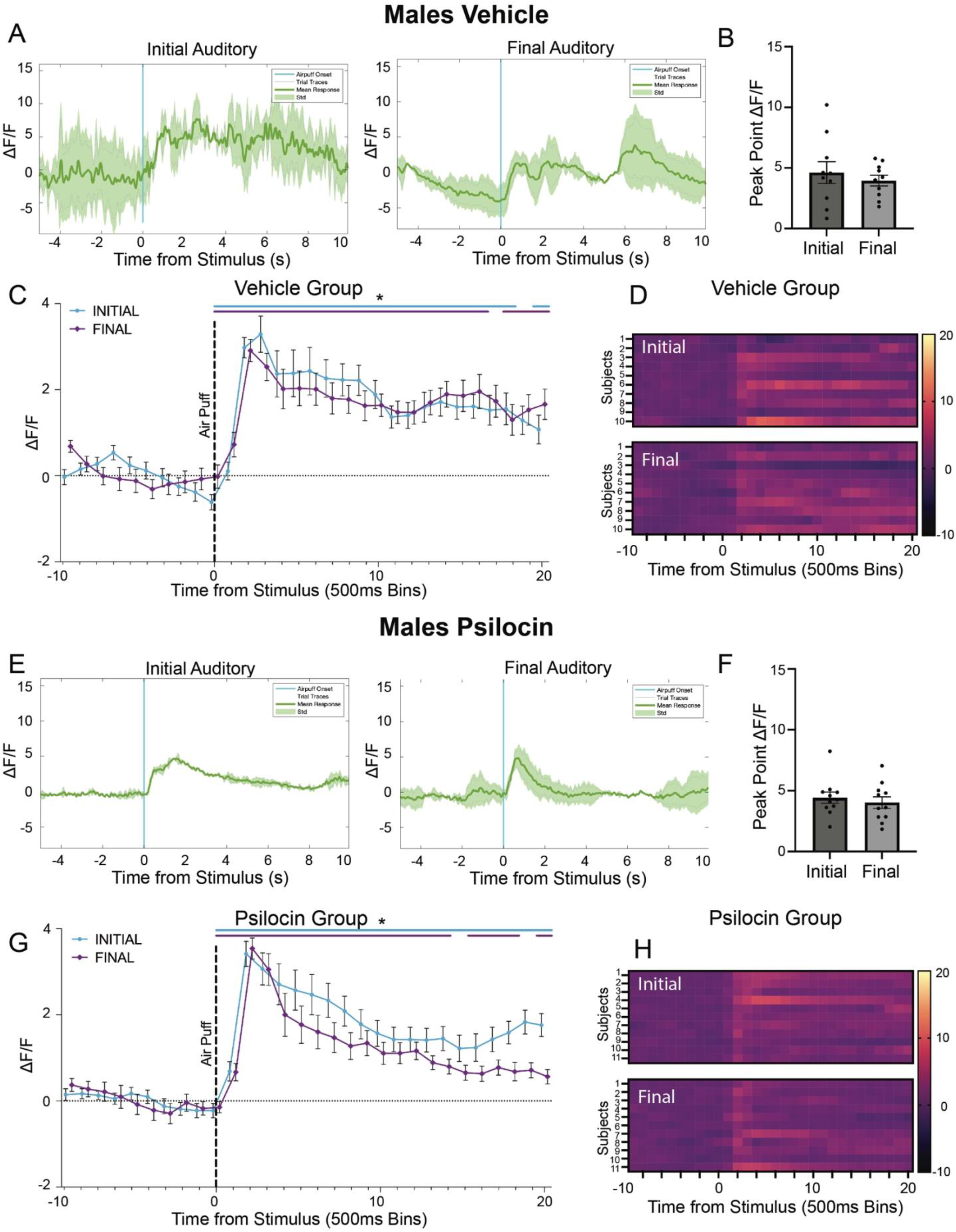
Changes in CeA Reactivity in Response to an Auditory Stimulus Following Psilocin Administration in Male Subjects. **(A)** Male vehicle group: representative mean ΔF/F traces showing an individual at initial auditory compared to final auditory. Blue line = air-puff onset, Grey lines = individual traces, Green line = mean trace, Std= standard deviation **(B)** Male vehicle group: peak point of binned mean ΔF/F traces compared between initial and final auditory recordings. Data points represent each individual subject’s peak ΔF/F value +/− SEM. **(C)** Male vehicle group: air-puff trace plots of changes in CeA fluorescence following exposure to a 500ms air-puff at 85 psi. Data points represent group averages within 500ms binned window +/− S.E.M. **(D)** Male vehicle group: Heatmaps comparing initial to final auditory. Each row represents an individual subjects mean trace. **(E)** Male psilocin group: representative mean ΔF/F traces showing an individual at initial auditory compared to final auditory. Blue line = air-puff onset, Grey lines = individual traces, Green line = mean trace, Std= standard deviation **(F)** Male psilocin group: peak point of binned mean ΔF/F traces compared between initial and final auditory recordings. Data points represent individual subject’s peak ΔF/F value +/− SEM. **(G)** Male psilocin group: air-puff trace plots of changes in CeA fluorescence following exposure to a 500ms air-puff at 85 psi. Data points represent group averages within 500ms binned window +/− S.E.M. **(H)** Male psilocin group: Heatmaps comparing initial to final auditory. Each row represents an individual subjects mean trace. In each trace bin plot panel, a significant increase in ΔF/F was determined whenever the lower bound of the 99% CI was >0. These points of statistical significance are shown as colored lines above each ΔF/F curve with colors corresponding to the respective binned traces with a * above the lines. \**p*<0.05, \*\**p*<0.01, *** *p*<0.001, **** *p*<0.0001.

## DISCUSSION

The present study explored the effects of the psychedelic compound psilocin on CeA activity using an immunohistological activity marker and on CeA reactivity to an aversive stimulus using *in-vivo* fiber photometry recordings during acute drug exposure and at 2-, 6-, and 28-days post drug administration. The study examined multi-level neurobiological and behavioral effects of psilocin by looking at acute drug effects, long-term alterations in activity following drug administration, and by probing for potential sex-specific effects of both injection and time. Psilocin administration produced increases in c-Fos expression in both males and females and sub-region analysis showed that increases in the female CeC were significantly greater in psilocin compared to the vehicle control (**Fig. 2B**). CeA reactivity in response to the air-puff stimulus following psilocin administration also displayed sex- and time-specific effects. In females, psilocin increased CeA reactivity acutely as compared to vehicle controls (**Fig. 2E-H**). Interestingly, both vehicle and psilocin groups showed increases in reactivity at specific follow-up timepoints compared to baseline. In the vehicle control females, there were large increases in the duration of CeA response at the 2- and 28-day follow-up. In the psilocin treated females, there were small increases at the 2- and 28-day follow-up, with a much larger response seen at the 6-day follow-up. Overall, we interpret these data to suggest there is variability in female CeA reactivity as opposed to any effect of the drug. In contrast, the male psilocin group did not show the same increased acute reactivity seen in the females (**Fig. 2F, 2H**) and instead displayed decreases in CeA reactivity at 2-day, 6-day, and 28-day follow-up recordings (**Fig. 3F, 3H**) that were not seen in vehicle control males (**Fig. 3G**). These long-term reductions in reactivity seen in males could be due to plasticity dependent changes in innervating circuitry. Future studies will investigate a potential upstream mediator of these observed effects.

In this study, we have replicated previous work demonstrating an increase in basal activity following administration of a 5-HT_2AR_ agonist psychedelic [37]. This could in part be due to local 5-HT_2AR_ excitatory activation in the CeA. However, the CeA contains a high proportion of 5-HT_1AR_, a presynaptic receptor with inhibitory mechanisms [59]. Another potential explanation lies in more of a circuit/systems level effect of the drug. For instance, the dorsal raphe nucleus is a region within the brainstem that is involved in serotonin synthesis that also has a high density of 5-HT_1AR_. Previous work has shown that administration of psychedelics produces a local decrease in activity within the DRN that is likely driven by 5-HT_1AR_ [60]. Additionally, it has been shown that there are dense projections of serotonergic neurons from the DRN to the locus coeruleus (LC) [61], and that serotonin release within the LC produces a tonic inhibition of LC neurons [62]. Thus, given previously reported noradrenergic projections from LC to CeA [63], it is possible that disinhibition of the LC through inhibition of the DRN via activation of DRN 5-HT_1ARs_ could result in increased basal activity within the CeA. Future work will look at potential circuit-based mediators of both the observed changes in basal activity and changes in stimulus reactivity within the CeA.

Clinical studies have illustrated that psilocybin produces alterations in amygdala reactivity and connectivity [8, 12, 15, 17, 39–41]. Here, we utilized a preclinical approach to examine the actions of psilocin, the psychoactive metabolite of psilocybin, on CeA activity in a rodent model. As most clinical studies report a reduction in amygdala reactivity or connectivity that correlated with therapeutic improvements [8, 15, 17, 39–41], we hypothesized that psilocin would produce similar decreases in reactivity in our model. Here we found that a single dose of psilocin produced persistent reductions in CeA reactivity to an aversive stimulus in male but not female rats. These findings lend to recent studies in both clinical [11, 13, 64] and preclinical [65] literature showing persistent reductions in depressive behavior after a single dose of psilocybin. However, given the increases seen in the female group, these data do not perfectly align with clinical findings with samples that include representation from female participants. One potential explanation could be that human fMRI lacks the resolution needed to truly parse apart amygdalar sub-nuclei activation. It is possible that effects seen in the CeA would not match effects seen in the basolateral amygdala (BLA). Additionally, the amygdala has been shown to be involved in both positive and negative valence, and therefore functioning within this region seems to be very dynamic and stimulus-dependent. A surprise air puff to a rat may be an entirely different experience subjectively than seeing an emotional face, and therefore could result in far different functioning within the amygdala complex. Additional experiments comparing drug effects on whole brain connectivity/activity will be addressed in future studies. While the specific mechanism behind these therapeutic effects remains unknown, this study provides evidence that changes in CeA activity may play a role in observed therapeutic effects.

Lateralization of amygdala response to fear processing has been noted in both preclinical [66] and clinical literature [67, 68], however there are contradictions in these data and therefore no clear, resolute consensus. For instance, one study in human imaging showed that males had greater activation in the right amygdala as compared to females during an emotional face perception fMRI task utilizing angry facial expressions [67]. Another human imaging study employing a similar task showed that amygdala activation differed based on the valence of the stimuli. Additionally, while males had more laterality, both male and female subjects showed greater activation in the left amygdala to fearful faces [68]. This is particularly relevant to the current study given that recordings were taken from a single hemisphere. Initially, hemispheres were chosen based on consistency and quality of signal. Within each group for both males and females, there is representation from both left and right hemispheres. While this does offer some protection from potentially confounding effects of amygdala lateralization, future work will be conducted to determine if there is an effect of amygdala lateralization on psilocin-induced changes in reactivity.

We report sex-specific effects of psilocin on basal activity and stimulus-induced reactivity within the CeA. Changes in basal activity were measured by c-Fos, an immediate early gene shown to be upregulated following neuronal activation [69]. We observed an increase in CeA c-Fos expression in both males and females following administration of psilocin, with region specific increases found in the CeC subregion in females. CeA reactivity was assessed using fiber photometry to measure in-vivo stimulus-induced fluctuations in calcium signaling. In the female psilocin group, we observed an acute increase in reactivity to the air-puff stimulus while on the drug that was not seen in the males. Additionally, we saw increases in reactivity at all of the follow-up recording days, while we saw reductions in CeA reactivity in the males on follow-up recording days (**Fig. 3B, 3D, 3E, 3G**). A recent study looking at cerebral blood flow and the expression of the serotonin transporter (5-HTT) showed that, in response to an aversive predator odor, there were increases in blood flow and c-Fos expression in the CeA that were related to 5-HTT expression in males but not females [70]. These findings suggest potential sex-specific expression of serotonin receptors within the amygdala. Given the affinity of psilocin for the serotonin system, differential expression of serotonin receptors within the CeA or surrounding nuclei could explain the sex-specific effects seen in this study. Additionally, a previous study showed sex-specific differences in 5-HT_2AR_ expression at baseline, wherein females had lower expression in the orbitofrontal cortex (OFC) than males [71]. Another potential explanation for the differential effects of psilocin between males and females may be due to differences in CeA circuitry. For example, the OFC has been shown, in rodents and non-human primates, to have direct connections to the CeA as well as to the intercalated nuclei, a local GABAergic population within the amygdala complex [72, 73]. Lack of activation of this circuit due to a reduced expression of OFC 5-HT_2AR_ in females could prevent an inhibition of CeA firing, thus causing the increase in reactivity seen while on psilocin in females but not males. Future experiments are needed to examine the effects of psilocin on CeA circuits innervating distinct brain regions.

In the presence of a threatening stimulus, animals will employ either active or passive coping strategies. In this study, the active threat response was characterized by darting, or immediate fleeing from the air-puff stimulus, while the passive threat response was characterized by immobility following the air-puff. Here, we show that females predominantly exhibited an active response, while males were more evenly split between active and passive responses. In male psilocin-treated animals who exhibited an active response at baseline, there was a significant decrease in the amplitude of CeA reactivity following exposure to the air-puff at the 2-day follow-up, with no changes in vehicle control. This decrease in reactivity was not seen in the passive-responder vehicle or psilocin groups. While further interpretation regarding adaptive vs. maladaptive strategies in this context is outside the scope of the current study, these results suggest that there may be differences in CeA reactivity and threat responding strategy that are implicated in the efficacy of psychedelics.

To determine the effects of psilocin on associative learning, we utilized an auditory-only stimulus recording taken after the initial baseline recordings and then again after the 28-day recording. Interestingly, in females the vehicle control group displayed reductions in CeA response magnitude to the auditory-only stimulus while reactivity in the males did not change. In the psilocin group however, reductions in CeA reactivity to the auditory stimulus were not observed in males or females (**Figs. 5, 6**). Compared to the changes in reactivity seen with the standard air-puff stimulus, the changes in reactivity to the auditory stimulus were contradictory in females and absent in males. These data suggest a more dynamic alteration in CeA reactivity that is stimulus specific. One potential interpretation of the sustained activation in the psilocin females vs. vehicle control is that psilocin may influence associative learning to an aversive stimulus. Several studies have shown that 5-HT_2A_ agonist psychedelics produce improvements in various dimensions of cognition, including associative learning [74–76]. Another potential interpretation of these findings is that psilocin created a more robust/meaningful memory of the air-puff. Many of the clinical studies cite participants rating their experience during a psilocybin treatment as one of the most meaningful experiences of their life [13, 77, 78]. Additionally, increases in plasticity could contribute to differences in memory reconsolidation during air-puff administration while on the drug.

Our findings demonstrate that the psychedelic compound, psilocin, produces dynamic, sex-specific changes in the CeA, a key nucleus within the amygdala complex. Here we showed that administration of psilocin produces increases in CeA c-Fos expression and that, in females, these increases were primarily driven by increases in expression in the CeC. We have also shown that there are sex-specific changes in CeA reactivity while on psilocin. Specifically, we show that a single administration of psilocin produced acute increases in CeA reactivity in females, but not males. Additionally, we show that persistent reductions in CeA reactivity are seen as early as 2 and as long as 28 days following administration of the drug in males, but not females. Additionally, we show that these reductions in reactivity within the male CeA seem to be driven by those subjects that employed an active coping strategy to the air puff stimulus at baseline, suggesting that whatever neurobiological mechanism underlies these behaviors may also contribute to the long-term effects of psilocin. Given that dysregulation of the amygdala is a hallmark in many different psychiatric disorders, including anxiety, MDD, and PTSD [29, 31–36] and that psychedelic compounds show promise in treating many of these illnesses [4, 6–11], these findings provide important information on subregion-specific alterations in CeA function that may play a part in observed therapeutic effects of psychedelics. One limitation of this work is that no behavioral indices of hallucination (head twitch, wet back shakes, etc.) were collected following administration of the drug. Therefore, no claims can be made regarding the effects of hallucinations on amygdala activity following psychedelic drug exposure. Additionally, without the use of antagonists or known 5-HT_2AR_ mediated behavioral effects, we cannot claim any sort of receptor specificity in our findings. Though we are unable to implicate involvement of 5-HT_2AR_ in observed effects, it is important to note that while psilocin is an agonist at the 5-HT_2AR_, it is also a potent agonist at a variety of other serotonin receptors [79]. Future work will be required to explore cell-type/receptor-type specificity as well as in probing potential circuit-based mediators contributing to these observed alterations in CeA reactivity. However, the current findings provide important evidence for CeA alterations underlying the potential therapeutic effects of psilocin and psilocybin.

## ACKNOWLEDGEMENTS

This work was supported by National Institute of Health grants AA026858 (M.A.H.), GM135095 (D.P.E.), AA007573 (S.G.Q.), and by the Brain and Behavior Research Foundation NARSAD Young Investigator Award (M.A.H.)

## REFERENCES

1. Nichols, D.E., Hallucinogens. Pharmacology & therapeutics, 2004. 101(2): p. 131–181.

2. Dinis-Oliveira, R.J., Metabolism of psilocybin and psilocin: clinical and forensic toxicological relevance. Drug metabolism reviews, 2017. 49(1): p. 84–91.

3. Passie, T., et al., The pharmacology of psilocybin. Addiction biology, 2002. 7(4): p. 357–364.

4. Davis, A.K., et al., Effects of psilocybin-assisted therapy on major depressive disorder: a randomized clinical trial. JAMA psychiatry, 2021. 78(5): p. 481–489.

5. Bogenschutz, M.P. and M.W. Johnson, Classic hallucinogens in the treatment of addictions. Progress in Neuro-Psychopharmacology and Biological Psychiatry, 2016. 64: p. 250–258.

6. Johnson, M.W., A. Garcia-Romeu, and R.R. Griffiths, Long-term follow-up of psilocybin-facilitated smoking cessation. The American journal of drug and alcohol abuse, 2017. 43(1): p. 55–60.

7. Johnson, M.W. and R.R. Griffiths, Potential therapeutic effects of psilocybin. Neurotherapeutics, 2017. 14(3): p. 734–740.

8. Carhart-Harris, R.L., et al., Psilocybin for treatment-resistant depression: fMRI-measured brain mechanisms. Scientific reports, 2017. 7(1): p. 1–11.

9. Griffiths, R.R., et al., Psilocybin produces substantial and sustained decreases in depression and anxiety in patients with life-threatening cancer: A randomized double-blind trial. Journal of psychopharmacology, 2016. 30(12): p. 1181–1197.

10. Bogenschutz, M.P., et al., Psilocybin-assisted treatment for alcohol dependence: a proof-of-concept study. Journal of psychopharmacology, 2015. 29(3): p. 289–299.

11. Ross, S., et al., Rapid and sustained symptom reduction following psilocybin treatment for anxiety and depression in patients with life-threatening cancer: a randomized controlled trial. Journal of psychopharmacology, 2016. 30(12): p. 1165–1180.

12. Roseman, L., et al., Increased amygdala responses to emotional faces after psilocybin for treatment-resistant depression. Neuropharmacology, 2018. 142: p. 263–269.

13. Agin-Liebes, G.I., et al., Long-term follow-up of psilocybin-assisted psychotherapy for psychiatric and existential distress in patients with life-threatening cancer. Journal of Psychopharmacology, 2020. 34(2): p. 155–166.

14. Barrett, F.S., et al., Psilocybin acutely alters the functional connectivity of the claustrum with brain networks that support perception, memory, and attention. NeuroImage, 2020. 218: p. 116980.

15. Grimm, O., et al., Psilocybin modulates functional connectivity of the amygdala during emotional face discrimination. Eur Neuropsychopharmacol, 2018. 28(6): p. 691–700 DOI: 10.1016/j.euroneuro.2018.03.016.

16. Preller, K.H., et al., Psilocybin induces time-dependent changes in global functional connectivity. Biological psychiatry, 2020. 88(2): p. 197–207.

17. Mertens, L.J., et al., Therapeutic mechanisms of psilocybin: changes in amygdala and prefrontal functional connectivity during emotional processing after psilocybin for treatment-resistant depression. Journal of Psychopharmacology, 2020. 34(2): p. 167–180.

18. Roseman, L., et al., The effects of psilocybin and MDMA on between-network resting state functional connectivity in healthy volunteers. Frontiers in human neuroscience, 2014. 8: p. 204.

19. Carhart-Harris, R.L., et al., Neural correlates of the psychedelic state as determined by fMRI studies with psilocybin. Proceedings of the National Academy of Sciences, 2012. 109(6): p. 2138–2143.

20. Shao, L.-X., et al., Psilocybin induces rapid and persistent growth of dendritic spines in frontal cortex in vivo. Neuron, 2021. 109(16): p. 2535–2544. e4.

21. Jones, K.A., et al., Rapid modulation of spine morphology by the 5-HT2A serotonin receptor through kalirin-7 signaling. Proceedings of the National Academy of Sciences, 2009. 106(46): p. 19575–19580.

22. Ly, C., et al., Psychedelics promote structural and functional neural plasticity. Cell reports, 2018. 23(11): p. 3170–3182.

23. Lee, H.-M. and B.L. Roth, Hallucinogen actions on human brain revealed. Proceedings of the National Academy of Sciences, 2012. 109(6): p. 1820–1821.

24. Kometer, M., et al., Activation of serotonin 2A receptors underlies the psilocybin-induced effects on α oscillations, N170 visual-evoked potentials, and visual hallucinations. Journal of Neuroscience, 2013. 33(25): p. 10544–10551.

25. Kometer, M., et al., Psilocybin biases facial recognition, goal-directed behavior, and mood state toward positive relative to negative emotions through different serotonergic subreceptors. Biological psychiatry, 2012. 72(11): p. 898–906.

26. Halberstadt, A.L., et al., Differential contributions of serotonin receptors to the behavioral effects of indoleamine hallucinogens in mice. Journal of psychopharmacology, 2011. 25(11): p. 1548–1561.

27. Halberstadt, A.L., S.B. Powell, and M.A. Geyer, Role of the 5-HT2A receptor in the locomotor hyperactivity produced by phenylalkylamine hallucinogens in mice. Neuropharmacology, 2013. 70: p. 218–227.

28. Madsen, M.K., et al., Psychedelic effects of psilocybin correlate with serotonin 2A receptor occupancy and plasma psilocin levels. Neuropsychopharmacology, 2019. 44(7): p. 1328–1334 DOI: 10.1038/s41386-019-0324-9.

29. Gilpin, N.W., M.A. Herman, and M. Roberto, The central amygdala as an integrative hub for anxiety and alcohol use disorders. Biological psychiatry, 2015. 77(10): p. 859–869.

30. Ciocchi, S., et al., Encoding of conditioned fear in central amygdala inhibitory circuits. Nature, 2010. 468(7321): p. 277–282.

31. Sandu, A.L., et al., Amygdala and regional volumes in treatment-resistant versus nontreatment-resistant depression patients. Depression and anxiety, 2017. 34(11): p. 1065–1071.

32. Armony, J.L., et al., Amygdala response in patients with acute PTSD to masked and unmasked emotional facial expressions. American Journal of Psychiatry, 2005. 162(10): p. 1961–1963.

33. Frodl, T., et al., Enlargement of the amygdala in patients with a first episode of major depression. Biological psychiatry, 2002. 51(9): p. 708–714.

34. Rauch, S.L., et al., Exaggerated amygdala response to masked facial stimuli in posttraumatic stress disorder: a functional MRI study. Biological psychiatry, 2000. 47(9): p. 769–776.

35. Semple, W.E., et al., Higher brain blood flow at amygdala and lower frontal cortex blood flow in PTSD patients with comorbid cocaine and alcohol abuse compared with normals. Psychiatry, 2000. 63(1): p. 65–74.

36. Siegle, G.J., et al., Increased Amygdala and Decreased Dorsolateral Prefrontal BOLD Responses in Unipolar Depression: Related and Independent Features. Biological Psychiatry, 2007. 61(2): p. 198–209 DOI: https://doi.org/10.1016/j.biopsych.2006.05.048.

37. Gresch, P., L. Strickland, and E. Sanders-Bush, Lysergic acid diethylamide-induced Fos expression in rat brain: role of serotonin-2A receptors. Neuroscience, 2002. 114(3): p. 707–713.

38. Pędzich, B.D., et al., Effects of a psychedelic 5-HT2A receptor agonist on anxiety-related behavior and fear processing in mice. Neuropsychopharmacology, 2022: p. 1–11.

39. Kraehenmann, R., et al., Psilocybin-Induced Decrease in Amygdala Reactivity Correlates with Enhanced Positive Mood in Healthy Volunteers. Biol Psychiatry, 2015. 78(8): p. 572–81 DOI: 10.1016/j.biopsych.2014.04.010.

40. Kraehenmann, R., et al., The mixed serotonin receptor agonist psilocybin reduces threat-induced modulation of amygdala connectivity. NeuroImage: Clinical, 2016. 11: p. 53–60.

41. Barrett, F.S., et al., Emotions and brain function are altered up to one month after a single high dose of psilocybin. Scientific Reports, 2020. 10(1): p. 1–14.

42. Bas-Hoogendam, J.M., et al., Amygdala hyperreactivity to faces conditioned with a social-evaluative meaning–a multiplex, multigenerational fMRI study on social anxiety endophenotypes. NeuroImage: Clinical, 2020. 26: p. 102247.

43. Donegan, N.H., et al., Amygdala hyperreactivity in borderline personality disorder: implications for emotional dysregulation. Biological psychiatry, 2003. 54(11): p. 1284–1293.

44. Yang, T.T., et al., Adolescents with major depression demonstrate increased amygdala activation. Journal of the American Academy of Child & Adolescent Psychiatry, 2010. 49(1): p. 42–51.

45. Joos, A.A., et al., Amygdala hyperreactivity in restrictive anorexia nervosa. Psychiatry Research: Neuroimaging, 2011. 191(3): p. 189–195.

46. Moriarty, O., et al., Validation of an air-puff passive-avoidance paradigm for assessment of aversive learning and memory in rat models of chronic pain. Journal of neuroscience methods, 2012. 204(1): p. 1–8.

47. Veinante, P., I. Yalcin, and M. Barrot, The amygdala between sensation and affect: a role in pain. Journal of molecular psychiatry, 2013. 1(1): p. 1–14.

48. Wilson, T.D., et al., Dual and opposing functions of the central amygdala in the modulation of pain. Cell reports, 2019. 29(2): p. 332–346. e5.

49. Horsley, R.R., et al., Psilocin and ketamine microdosing: effects of subchronic intermittent microdoses in the elevated plus-maze in male Wistar rats. Behavioural pharmacology, 2018. 29(6): p. 530–536.

50. Spain, A., et al., Neurovascular and neuroimaging effects of the hallucinogenic serotonin receptor agonist psilocin in the rat brain. Neuropharmacology, 2015. 99: p. 210–220.

51. Barker, D.J., et al., Lateral preoptic control of the lateral habenula through convergent glutamate and GABA transmission. Cell reports, 2017. 21(7): p. 1757–1769.

52. Lerner, T.N., et al., Intact-brain analyses reveal distinct information carried by SNc dopamine subcircuits. Cell, 2015. 162(3): p. 635–647.

53. Friard, O. and M. Gamba, BORIS: a free, versatile open-source event-logging software for video/audio coding and live observations. Methods in ecology and evolution, 2016. 7(11): p. 1325–1330.

54. Schneider, C.A., W.S. Rasband, and K.W. Eliceiri, NIH Image to ImageJ: 25 years of image analysis. Nature methods, 2012. 9(7): p. 671–675.

55. Bland, J.M. and D.G. Altman, Statistics notes: bootstrap resampling methods. bmj, 2015. 350.

56. Bird, K.D., Analysis of variance via confidence intervals. 2004: Sage.

57. Choi, E.A., et al., Paraventricular thalamus controls behavior during motivational conflict. Journal of Neuroscience, 2019. 39(25): p. 4945–4958.

58. Gruene, T.M., et al., Sexually divergent expression of active and passive conditioned fear responses in rats. Elife, 2015. 4: p. e11352.

59. Albert, P.R. and F. Vahid-Ansari, The 5-HT1A receptor: signaling to behavior. Biochimie, 2019. 161: p. 34–45.

60. Sprouse, J.S. and G.K. Aghajanian, Electrophysiological responses of serotoninergic dorsal raphe neurons to 5-HT1A and 5-HT1B agonists. Synapse, 1987. 1(1): p. 3–9.

61. Morgane, P.J. and M. Jacobs, Raphe projections to the locus coeruleus in the rat. Brain research bulletin, 1979. 4(4): p. 519–534.

62. Kaehler, S.T., N. Singewald, and A. Philippu, Dependence of serotonin release in the locus coeruleus on dorsal raphe neuronal activity. Naunyn-Schmiedeberg’s archives of pharmacology, 1999. 359(5): p. 386–393.

63. Gu, Y., et al., A brainstem-central amygdala circuit underlies defensive responses to learned threats. Molecular psychiatry, 2020. 25(3): p. 640–654.

64. Carhart-Harris, R.L., et al., Psilocybin with psychological support for treatment-resistant depression: six-month follow-up. Psychopharmacology, 2018. 235(2): p. 399–408.

65. Hibicke, M., et al., Psychedelics, but not ketamine, produce persistent antidepressant-like effects in a rodent experimental system for the study of depression. ACS chemical neuroscience, 2020. 11(6): p. 864–871.

66. Baker, K.B. and J.J. Kim, Amygdalar lateralization in fear conditioning: evidence for greater involvement of the right amygdala. Behavioral neuroscience, 2004. 118(1): p. 15.

67. Schneider, S., et al., Boys do it the right way: sex-dependent amygdala lateralization during face processing in adolescents. Neuroimage, 2011. 56(3): p. 1847–1853.

68. Killgore, W.D. and D.A. Yurgelun-Todd, Sex differences in amygdala activation during the perception of facial affect. Neuroreport, 2001. 12(11): p. 2543–2547.

69. Morgan, J. and T. Curran, Calcium as a modulator of the immediate-early gene cascade in neurons. Cell calcium, 1988. 9(5-6): p. 303–311.

70. Kolter, J.F., et al., Serotonin transporter genotype modulates resting state and predator stress-induced amygdala perfusion in mice in a sex-dependent manner. PloS one, 2021. 16(2): p. e0247311.

71. Pitychoutis, P., et al., 5-HT1A, 5-HT2A, and 5-HT2C receptor mRNA modulation by antidepressant treatment in the chronic mild stress model of depression: sex differences exposed. Neuroscience, 2012. 210: p. 152–167.

72. REMPEL-CLOWER, N.L., Role of orbitofrontal cortex connections in emotion. Annals of the New York Academy of Sciences, 2007. 1121(1): p. 72–86.

73. Barbas, H., Flow of information for emotions through temporal and orbitofrontal pathways. Journal of anatomy, 2007. 211(2): p. 237–249.

74. Gimpl, M., I. Gormezano, and J. Harvey, Effects of LSD on learning as measured by classical conditioning of the rabbit nictitating membrane response. Journal of Pharmacology and Experimental Therapeutics, 1979. 208(2): p. 330–334.

75. Doss, M.K., et al., Psilocybin therapy increases cognitive and neural flexibility in patients with major depressive disorder. Translational psychiatry, 2021. 11(1): p. 1–10.

76. Roberts, M. and P. Bradley, Studies on the effects of drugs on performance of a delayed discrimination. Physiology & Behavior, 1967. 2(4): p. 389–397.

77. Griffiths, R.R., et al., Mystical-type experiences occasioned by psilocybin mediate the attribution of personal meaning and spiritual significance 14 months later. Journal of psychopharmacology, 2008. 22(6): p. 621–632.

78. Griffiths, R.R., et al., Psilocybin can occasion mystical-type experiences having substantial and sustained personal meaning and spiritual significance. Psychopharmacology, 2006. 187(3): p. 268–283.

79. PDSP Ki Database.

